# NEMO recruitment at single cytokine-receptor complexes shows quantized dynamics independent of ligand affinity

**DOI:** 10.1101/2025.04.18.649561

**Authors:** A. Hyun Kim, Benjamin Krummenacher, Jason Yeung, David R. Koes, Robin E. C. Lee

**Author notes:** Authors contributed equally (A. Hyun Kim, Benjamin Krummenacher).

## Abstract

Cells use a limited number of receptors to sense and process molecular information from their environment. In the classical view of signaling, receptor-ligand affinities determine binding kinetics, in timescales of diffusion, where their time-averaged contact duration regulates rapid cytoplasmic signaling events to coordinate cellular responses. For some cytokines, single receptor-ligand binding events can initiate large multiprotein complexes in the cytoplasm that assemble over tens of minutes, bringing to question how cytokine affinity influences the sensitivity and strength of signaling. Here, we leverage naturally occurring variation of IL-1β from multiple species to determine the impact of affinity on human IL-1 receptor signaling. Using experiments and models we investigate single receptor complexes activated by ligands that vary across multiple orders of magnitude in affinity. Our results show that while the receptor-ligand affinity establishes cytokine response sensitivity, activated IL-1 receptor complexes signal as discrete, quantized packets of signaling flux independent of affinity.

## Introduction

Cells rely on receptor-based signaling to interpret their molecular environment and regulate essential functions such as growth, differentiation, motility, adhesion, and immune responses^1^. Coordination of these diverse biological processes depends on binding between extracellular ligands and transmembrane receptors, which in turn regulate intracellular signaling cascades. In their classic paper, Berg and Purcell considered receptor-level mechanisms that cells use to accurately detect and respond to ligand concentrations in noisy, fluctuating molecular environments. Their theory explains how cells infer extracellular ligand concentrations by integrating the time-averaged occupancy of cell surface receptors, with an upper bound to signaling precision defined by molecular diffusion near the receptor^2^. More recent extensions of this work have incorporated fluctuation-dissipation theory to establish general limits of ligand-receptor signaling, demonstrating the importance of affinity in shaping the precision and accuracy of downstream responses^3–5^. At the core of these theories is the principle that the contact duration of a ligand with its receptor encodes downstream molecular signaling events. Consequently, a receptor’s affinity for its ligand is a central determinant of signaling dynamics through its impact on binding and unbinding kinetics.

Although originally developed to describe bacterial chemotaxis, these principles of ligand-receptor contact duration and affinity extend to many eukaryotic signaling systems, for example the superfamily of ligand-gated channels. Here, ion channel activation is allosterically regulated by the binding of a ligand to its receptor so contact duration directly controls its behavior^6^. Similarly, G protein-coupled receptor (GPCR) systems are well-established examples of how contact duration directly influences downstream signaling^1^. Acting as a molecular rheostat, ligand-bound GPCRs catalyze the exchange of GDP for GTP on the Gα subunit, triggering Gα dissociation from the Gβγ subunits and activating a multitude of downstream cascades^7^. The signal is terminated when the Gα subunit hydrolyzes its bound GTP, either spontaneously or with the help of GTPase-activating proteins. Because GPCRs can adopt multiple conformational states with varying activity levels, the strength and specificity of the signal depend on both ligand-receptor affinity and the dynamics of G-protein engagement^8^. For instance, the adenosine A1 receptor (A1R) is a GPCR that exhibits rapid signal termination upon ligand dissociation, leading to immediate deactivation of downstream effectors^9^. Similarly, dopamine D2 receptors (D2R) show prompt cessation of G-protein signaling upon ligand unbinding, emphasizing the critical role of ligand residence time in shaping cellular outcomes in disorders such as Parkinson’s disease and schizophrenia^10,11^.

Receptor tyrosine kinases (RTKs) also rely on ligand-binding duration to fine-tune intracellular signaling responses. Upon ligand engagement, RTKs dimerize and autophosphorylate, creating docking sites for adaptor proteins that recruit and activate effectors such as Ras, PI3K, and STATs^12^. The extent and duration of receptor phosphorylation—and the stability of downstream complexes—are directly influenced by how long a ligand remains bound. Brief interactions lead to transient activation, favoring fast-turnover pathways, whereas prolonged engagement sustains phosphorylation and enhances recruitment of scaffold proteins like Grb2 and Gab1, amplifying MAPK and PI3K signaling^13,14^. RTK internalization via recycling or degradation is also dependent on ligand occupancy time, adding another layer of temporal regulation. As with GPCRs, RTKs adjust their signaling output based on both ligand affinity and contact duration. Non-catalytic tyrosine-phosphorylated receptors (NTRs) are a similar class of receptors that lack an intrinsic kinase domain and instead rely on recruitment of Src-family kinases for phosphorylation. T-cell receptors are essential NTRs that orchestrate immunity through kinetic proofreading, where affinity-mediated contact duration of highly similar antigenic ligands leads to either weak and transient or explosive and sustained signaling^15,16^.

Dysregulation of receptor systems such as GPCRs and RTKs is frequently implicated in disease, where altered affinity or binding duration drives pathological cell behaviors. In cancer, aberrant RTK activation often results from elevated ligand affinity or receptor overexpression, resulting in sustained proliferative signaling and resistance to apoptosis^17–19^. Specifically, mutations to epidermal growth factor receptors (EGFRs) extracellular domains (e.g., R84K, A265V, G574V) significantly increase ligand-binding affinity^20^. These mutations heighten ligand potency in stimulating receptor autophosphorylation and downstream pathways including phospholipase Cγ, Akt, and MAP kinase, thereby contributing to oncogenic transformation^20^. Likewise, in neurodegenerative diseases, altered GPCR dynamics compromise neurotransmission. For example, Alzheimer’s Disease (AD) patients show reduced ligand binding to serotonin and adrenergic receptors, a deficit that contributes to cognitive impairment^21^. In both contexts, changes in receptor-ligand interaction kinetics, whether through altered affinity or contact duration, profoundly reshape cellular responses. These examples highlight how finely tuned physical parameters, such as residence time, play decisive roles in both health and disease.

The nuclear factor kB (NF-κB) signaling pathway is an established model for investigating how extracellular cytokine signals are interpreted by cells to regulate specific gene expression programs^22–24^. As a master regulator of immune and inflammatory responses, NF-κB controls diverse physiological processes, from pathogen defense to tissue homeostasis. Dysregulation of this pathway is implicated in a broad range of diseases, including rheumatoid arthritis, inflammatory bowel disease, and cancer, underscoring its central role in maintaining immune balance^25–29^. Under basal conditions, NF-κB dimers are sequestered in the cytoplasm by their inhibitor, IκB, preventing transcriptional activity^30–32^. Canonical NF-κB signaling is initiated when inflammatory cytokines such as TNF or IL-1β bind to their receptors at the plasma membrane, triggering the assembly of receptor-proximal complexes that activate the IκB kinase (IKK) complex. IKK then phosphorylates IκB, targeting it for proteasomal degradation and liberating NF-κB to translocate into the nucleus and drive gene expression^33–35^.

Although TNF, IL-1β, and related inflammatory factors utilize distinct receptors, their signaling mechanisms converge at several key steps. TNF binding to TNFR1 recruits the adaptor protein TRADD, as well as RIPK1, TRAF2/5, and cIAPs to form Complex I (CI), a membrane-associated signaling hub that nucleates formation of non-degradative linear and branched ubiquitin chains. These ubiquitin scaffolds then serve as platforms for recruiting TAB/TAK1 and IKK complexes, facilitating their kinase activation. Similarly, IL-1β binding to IL-1R1 and its co-receptor IL-1R3 initiates the recruitment of MyD88, forming a CI-like structure that assembles with IRAK kinases^36^. Activated IRAK1 interacts with TRAF6 to generate similar polyubiquitin structures as CI and subsequently coordinates downstream signaling by regulating phosphorylation-dependent IKK activity^37–39^. In contrast to simpler receptor systems, following ligand-receptor binding, CI and CI-like signaling requires the formation of dynamic supramolecular assemblies that can be observed in living cells through an EGFP-fusion of the IKK regulatory subunit NEMO^40–42^. Collectively, CI-like complexes regulate multi-step feedback loops that determine the amplitude, duration, and specificity of downstream signaling responses^42,43^.

Despite extensive characterization of NF-kB signaling, a fundamental question remains: how do receptor-ligand binding kinetics, affinity, and contact duration shape downstream inflammatory signals? Previous results have shown that across a range of concentrations, IL-1β-induced CI-like complexes exhibited remarkable similarity in intensity and size^42^, suggesting a possible departure from affinity- and duration-regulated signaling. Here, we take advantage of naturally occurring sequence variation to probe the affinity space of IL-1β and quantitatively characterize its effect on dynamics of single activated receptor complexes. Using phylogenetic and structural analyses of IL-1β homologs, we infer that IL-1β from different species will bind to human IL-1R with altered kinetics. Using purified IL-1β from a selection of mammals, we characterize responses of human cells expressing a reporter for CI-like structures and observe shifted dose response curves where ligand affinity alters potency. Using computational models and single particle tracking (SPT), we show that despite inferred affinity variation over several orders of magnitude between ligand-receptor complexes, the signal strength and temporal dynamics of single activated CI-like complexes are invariant. We conclude that while receptor-ligand affinity fine-tunes the thresholds for receptor activation, the aggregate of signaling flux from each activated CI-like complex is quantized and independent of receptor-ligand affinity.

## Results

### Naturally occurring sequence and structure variation in IL-1β is predicted to alter affinity for IL-1R1

To probe the molecular basis of competent IL-1R1-dependent signal transmission, we performed a cross-species multiple sequence alignment of IL-1β, focusing on residues implicated in receptor binding. Using a curated panel of twelve mammalian species spanning over 100 million years of evolutionary divergence, we compared full-length IL-1β protein sequences and constructed a distance tree based on sequence dissimilarity (Figure 1A). This analysis revealed appreciable sequence variation between species, with clear stratification by evolutionary proximity; for example, IL-1β from rodents clustered separately from primate or ungulate sequences.

**Fig. 1:**
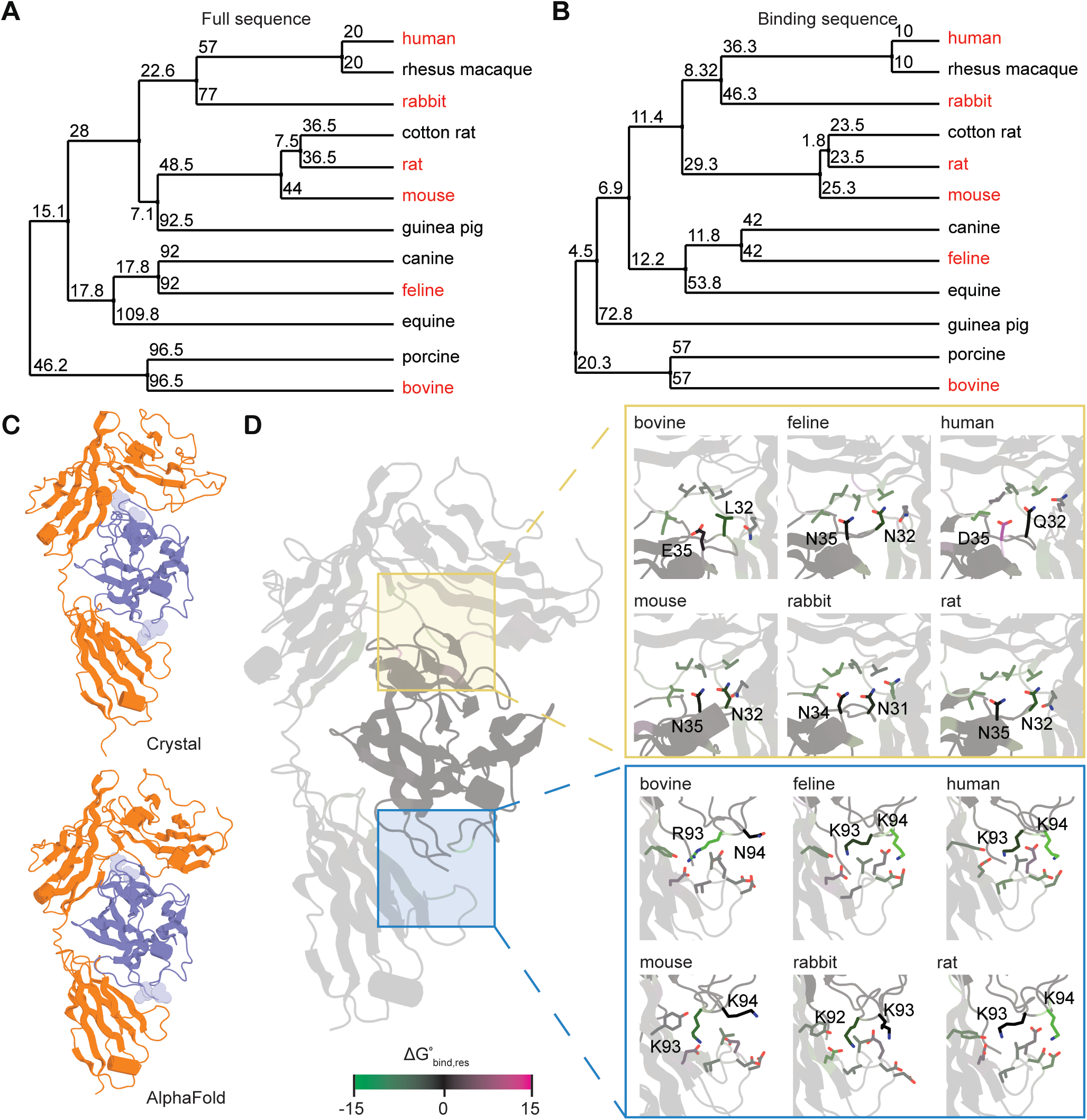
Cross-species comparison of IL-1β sequence and structure. **A.** Average BLOSUM62 distance trees across a panel of 12 species’ IL-1β full sequences. **B.** Average BLOSUM62 distance trees across a panel of 12 species’ IL-1β predicted binding regions only. Species in red font were selected for subsequent structural and experimental analysis. **C.** Crystal structure and generated structure for IL-1β (blue) in complex with the extracellular domain of IL-1R1 (orange). **D.** Differences in binding conformations across generated structures of 6 species’ IL-1β–IL-1R1 complexes for two regions heavily implicated in binding, with residues colored by their estimated contributions to the change in free energy.

We next restricted our comparison to the IL-1R1-binding interface by leveraging the human IL-1β–IL-1R1 co-crystal structure (PDB: 1ITB). From this structure, we identified residues directly contacting IL-1R1 and extracted the corresponding positions from each species’ sequence (Figure S1A). Interestingly, the dissimilarity tree based on these interface residues largely followed the hierarchy of the full-sequence tree, hinting at the possibility that natural variation at the binding interface may generally align with overall evolutionary divergence (Figure 1B). These findings raise the possibility that even subtle substitutions at receptor-contacting residues can modulate cytokine-receptor affinity and, by extension, downstream signaling responses in the cell. Thus, evolutionary drift in IL-1β’s binding interface may underpin receptor-ligand co-evolution or species-specific fine tuning for aspects of inflammatory signal transduction^44,45^.

Next, we assessed how sequence divergence at the IL-1R1-binding interface influences ligand–receptor affinity by simulating the dynamics of predicted chimeric IL-1β–IL-1R1 complexes across species. We first validated this approach by comparing AlphaFold predictions of the human IL-1β–IL-1R1 complex to its co-crystal structure^8^. The predicted complex closely recapitulated the experimental structure both in global architecture and in local binding conformations, supporting the method’s use for comparative analysis of this system (Figures 1C and S1B). For a selection of 5 species that were distributed along the dissimilarity distance tree, in addition to human, we then performed molecular dynamics simulations of IL-1β variants in complex with human IL-1R1. Analysis of predicted binding interfaces revealed varying degrees of conformational similarity to the human complex across species, particularly in loop regions previously implicated in receptor contact (Figure 1D). To quantify the effect of differences, we performed short simulations and calculated the total estimated binding free energy for each chimera using Molecular Mechanics/Generalized Born Surface Area (MM/GBSA)^46^. The differences in ΔG° values spanned nearly two orders of magnitude across the species panel, indicating a broad stratification of predicted binding affinity (Figure S1). The contribution to ΔG° of each residue was determined using the Molecular Mechanics/Poisson-Boltzmann Surface Area (MM/PBSA) method^46^. Regions of low ΔG° aligned with structurally determined binding interfaces and indicated local variation in binding favorability that may alter affinity (Figures 1 and S1).

Importantly, the predicted binding offered an independent metric of receptor engagement strength, which corresponds to the receptor-ligand binding affinity. For instance, the bovine IL-1β variant showed consistently higher predicted ΔG (weaker affinity) values compared to most other species and especially so for feline (strongest affinity), suggesting changes in affinity for the receptor even when overall sequence identity is moderately conserved. The predicted affinity for human IL-1β was lower than expectations, likely due to the generated conformation of Asp35, which shows an approximately four-fold increase in binding free energy relative to the crystal structure. However, this does not preclude the possibility that specific structural interactions of human IL-1β–IL-1R1 may allosterically facilitate other favorable interactions in the multiprotein CI-like receptor complex, despite reduced predicted affinity. Taken together, naturally occurring variation in IL-1β from feline to bovine gives rise to a spectrum of ligand-receptor conformations that can produce signaling-competent CI-like assemblies.

### CI-like Assemblies Observed in Living Cells Reveal Species-Specific Response Sensitivities

U2-OS expressing an endogenous fusion of EGFP-NEMO is a powerful tool to observe aspects of inflammatory receptor engagement because it provides two distinct outputs^42,43^. Over timescales of hours, it first informs when signalling competent CI-like receptor complexes form and dissipate based on the binary detection of NEMO assemblies. Second, from time courses of NEMO intensity at single assemblies in the timescales of seconds-to-minutes, cytokine-specific kinetics of NEMO recruitment and dissolution at each CI-like complex can be measured. Together, these observables at the plasma membrane are predictive of the sensitivity, timing, and strength of downstream NF-κB pathway activation in the same cell^42,43^.

To compare the dynamics of cytokine sensing across species, we examined the formation of CI-like structures by live-cell imaging of EGFP-NEMO in response to stimulation with recombinant IL-1β proteins from a selection of 6 species from our structural analysis (Figure 2; Movie S1). These cytoplasmic EGFP-NEMO assemblies represent upstream organizing hubs of the IKK complex, which form rapidly following cytokine stimulation and contribute to IKK enzymatic activation^40,42,43^. Using time-lapse imaging, we directly observed the assembly and disassembly of CI-like complexes, enabling us to measure signaling competency of the cytokines in single cells (Figures 2A and S2A). Remarkably, EGFP-NEMO at mature CI-like complexes following stimulation with IL-1β from all species were similar in appearance and intensity (Figures 2B and S2B), suggesting a conserved molecular composition once complexes were formed.

**Fig. 2:**
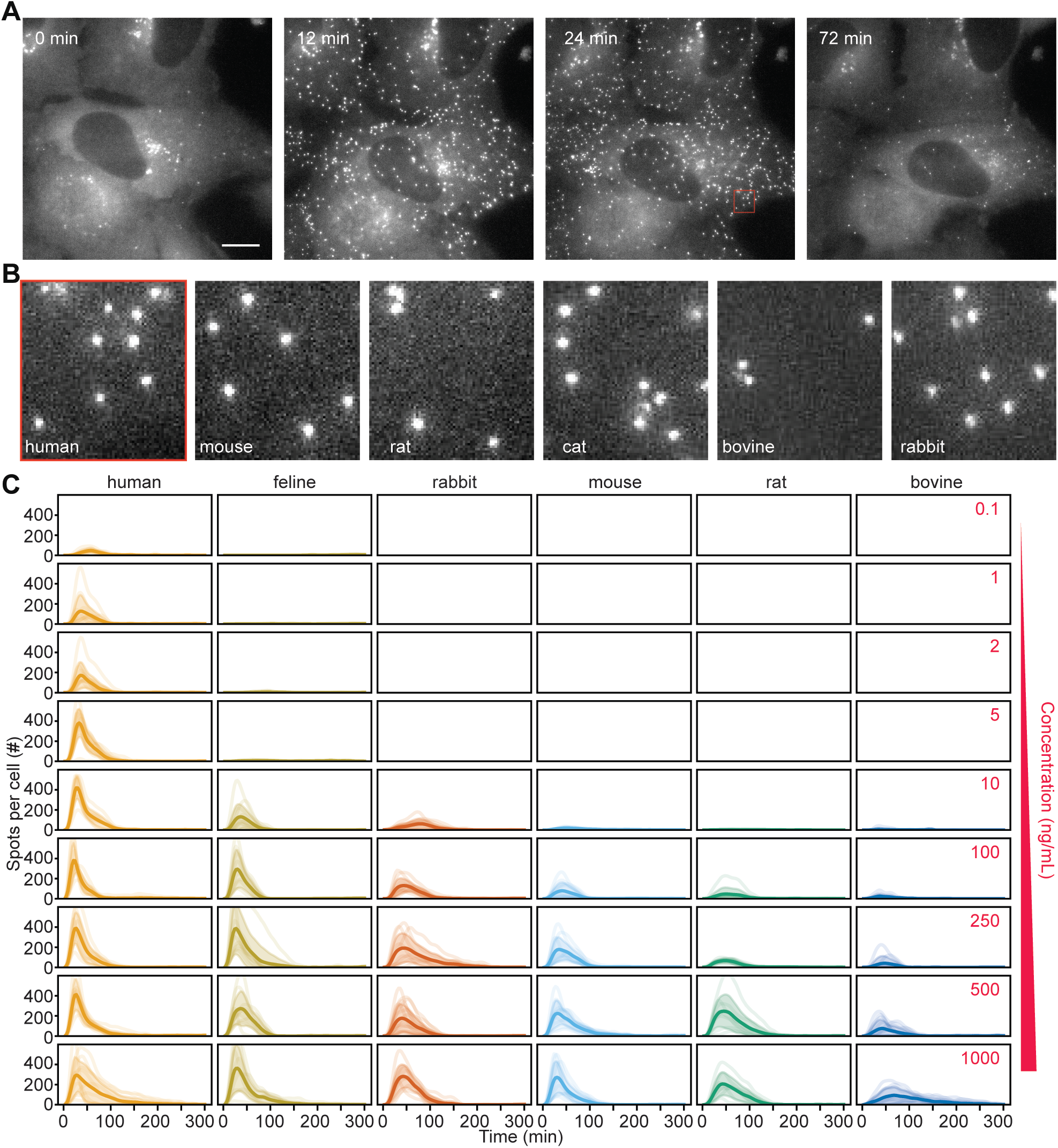
EGFP-NEMO assemblies form with varying sensitivities to IL-1β from different species. **A.** Time-lapse images of EGFP-NEMO endogenously expressed in U2OS cells stimulated with 250 ng/mL Human IL-1β. Scale bar, 20 µm. **B.** Magnification of EGFP-NEMO assemblies following 24 minutes of stimulation with 250 ng/mL IL-1β from indicated species. For scale, the red inset box is displayed in panel A. See also Movie S1 and Figure S2. **C.** Single-cell time courses for the number of EGFP-NEMO complexes in response to indicated concentrations and species of IL-1β.

Next, we measured EGFP-NEMO puncta across 4-orders of magnitude of IL-1β concentration for each species and extracted time course responses of IKK spot numbers per cell (Figure 2C). As previously, the number of EGFP-NEMO puncta peak within 30 minutes and typically adapt back to baseline values within 60-120 minutes. Comparing between species at the same concentration, the number of NEMO puncta varied substantially, reflecting differing levels of sensitivity. While all species showed stimulus-induced formation of NEMO puncta, the strength of the response in terms of both amplitude and duration varied considerably between species, with human showing the greatest sensitivity to lower concentrations, followed by feline as well as rabbits and rodents. Sensitivity of the response to bovine IL-1β was markedly lower than all other species. Moreover, response sensitivity to IL-1β from across species otherwise show the same general ranking as predicted by ΔG values (Figure S1D). These differences suggest that despite species-specific variability in IL-1β structure, all can produce signaling-competent CI-like complexes with activation thresholds that are consistent with their respective predicted affinities.

### Experiments and Models Show that Species-specific Ligand Affinities Span Over Log Decades

As described earlier, CI and CI-like complexes form sequentially from primary ligand-receptor binding, followed by recruitment of co-receptors and cytoplasmic adapter proteins before forming a mature signaling-competent complex (Figure 3A). We previously showed that features such as the “Area Under the Curve” (AUC) for the timing and number of CI-like complexes can be extracted from single-cell time courses of EGFP-NEMO and are strong predictors of dose-response characteristics^42^. Therefore, each trajectory was decomposed into a set of quantitative descriptors, including area under the curve (AUC), peak amplitude (Max), response timing (T_MAX_, FWHM, Adaptation Time), and rates of rise and fall (Rate_up_, Rate_down_; Figure 3B). These features capture both the strength and shape of the response, allowing us to assess the signaling competency and fidelity across species.

**Fig. 3:**
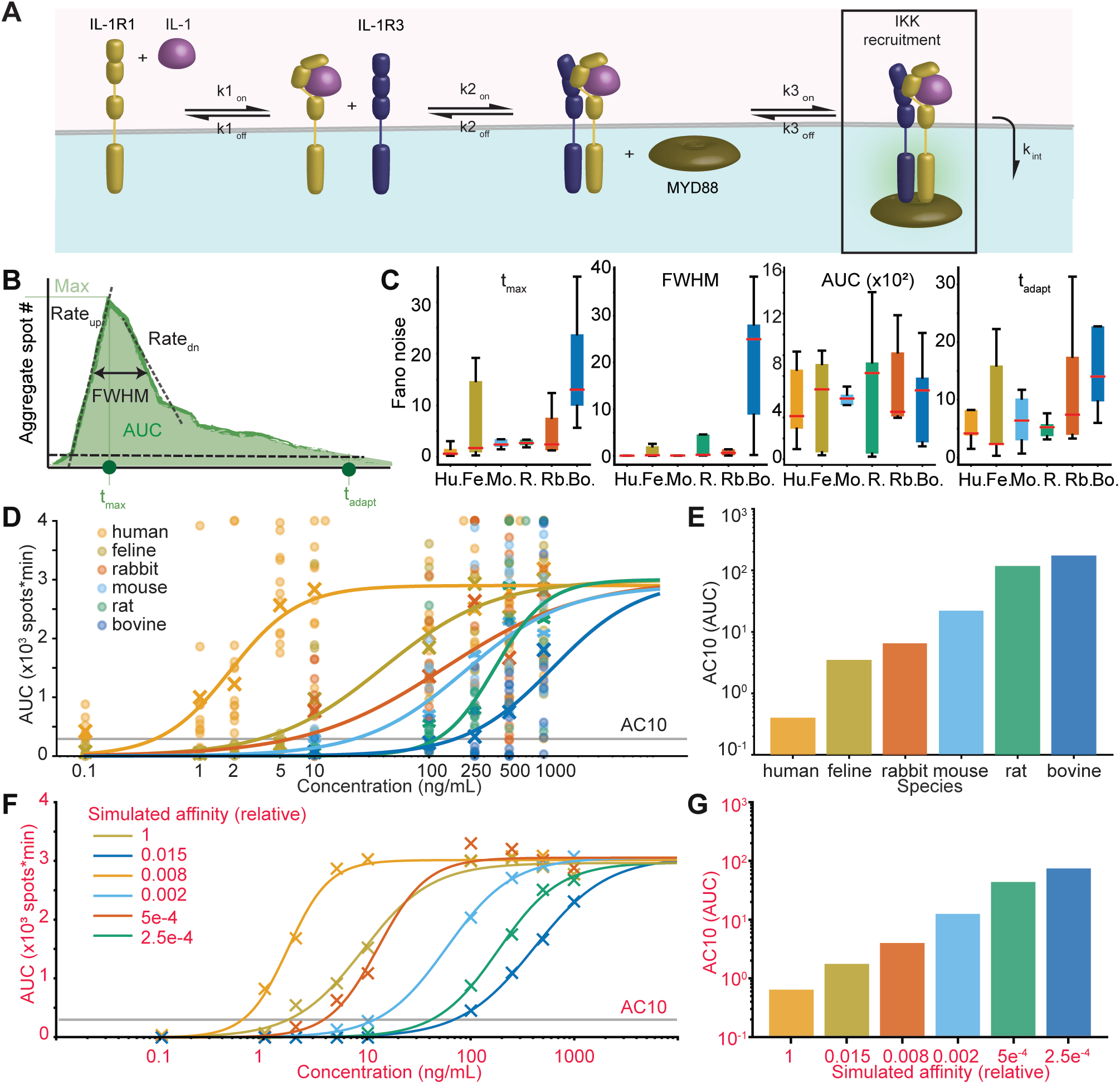
Dose responses of EGFP-NEMO dynamics predict the cross-species aggregate affinity for CI-like assemblies. **A.** Schematic of IL-1β-induced EGFP-NEMO complex formation at the plasma membrane. IL-1β first binds to IL-1R1, enabling recruitment of IL-1R3 to form the receptor complex. Cytoplasmic MYD88 then associates with the complex, facilitating IKK recruitment and the formation of EGFP-NEMO puncta. **B.** Quantitative descriptors extracted from each single-cell time courses of EGFP-NEMO puncta. *Rate_up_* and *Rate_dn_* describe the accumulation and dissipation of puncta. Max describes the maximum number of observed puncta at time *t_max_*. Area Under the Curve (AUC), time of adaptation (*t_adapt_*), and Full Width Half Maximum (FWHM), are also extracted for each single cell time course. **C.** Box plots of noise, defined by the Fano factor, evaluated for single-cell time courses at each concentration, and compared as box plots for each indicated descriptor and species. Human consistently shows the lowest noise, while bovine exhibits the highest. **D.** Sigmoid curves fitted to the mean values of each experimental single-cell descriptor across IL-1β concentrations and species, revealing dose–response relationships. The AC_10,_ defined as the concentration at which 10% of the maximal response is reached, is indicated. See Table S1 for fit parameters. **E.** AC_10_ values for each species derived from sigmoid fits to the experimental AUC descriptor, reflecting the concentration at which 10% of the maximal response is reached. **F.** Stochastic simulations using a minimal model recapitulate experimental results. Simulated dose–response curves as in (D), revealing dose–response relationships for each predicted affinity. See Table S2 for simulation parameters and Table S3 for fit parameters. **G.** AC_10_ values for each species derived from sigmoid fits to the simulated AUC descriptor, reflecting the predicted sensitivity to IL-1β. See Table S1B for fit parameters.

When comparing between species, we immediately noticed that there are differences in cell-to-cell variability that can be quantified by the Fano noise for each descriptor (Figure 3C). Notably, many descriptors showed the lowest Fano noise for human responses, followed by feline, rodents and rabbit, and the greatest variability in response to bovine IL-1β, suggesting that receptor engagement by cytokines from other species is less coordinated and noisier. Descriptors for rise and fall, as well as the max number of spots, did not show any species-specific trends.

Next, we measured dose responses for the AUC and max number of peaks, continuous dose response descriptors of EGFP-NEMO puncta that were previously shown to encode dosage information^42^. For each species, AUC and Max versus concentration were well fit by sigmoid curves again showing dose-response stratification from human to feline, rabbit and rodents, and bovine (Figures 3D and S3A; Table S1). To quantify sensitivity for each species, we calculated the concentration required to reach 10% of the maximal observed response (AC10; grey lines in Figures 3D and S3A) from sigmoid fits. For both descriptors, AC10 values spanned nearly three orders of magnitude between human and bovine IL-1β, where feline through rabbit and rodents were distributed in between (Figures 3E and S3B). This stratification further supports structural predictions for affinity, and the notion that human-human cytokine signaling is both more sensitive and robust.

To approximate how primary receptor–ligand binding affinity might shape the observed variation in signaling dynamics across species, we implemented a stochastic mechanistic model using the Gillespie method^47^. This minimal model captures the sequential assembly of CI-like complexes as a three-step reversible binding cascade: (1) IL-1β binds IL-1R1 to form the primary receptor–ligand complex, (2) the accessory protein joins this complex to form a trimeric intermediate, and (3) MYD88 binds as a terminal adaptor to complete the complex and initiate downstream NEMO recruitment (Figure 3A and STAR Methods). Once the binding cascade is complete, the complex dissociates either due to degradation or internalization^48,49^. Importantly, to isolate the contribution of primary binding affinity, we held all other kinetic parameters constant across species and varied only the on-rate and off-rate of the initial IL-1β–IL-1R1 interaction. The relative contribution of other components to forming the mature CI-like complex are relative to the primary binding. Hence, the simulated affinity of the IL-1β– IL-1R1 interaction is representative of the other interactions and therefore can be considered an “aggregate affinity” of IL-1β in its contribution to forming and binding the mature signaling-competent multi-protein complex. After fitting the model to human dose-responses for the number of CI-like complexes, we found that all other species could be recapitulated by scanning the aggregate affinity of IL-1β (Figure S3C; Table S2). Remarkably, by extracting AUC and MAX descriptors from simulated data, our results infer that the aggregate affinity of human IL-1β is 2 log decades stronger than feline and 4 log decades stronger than bovine, with rabbit and rodent affinities interspersed between (Figures 3 and S3; Table S3). These results suggest that the initial ligand–receptor interactions between IL-1R1 and IL-1β lead to significant changes to downstream kinetics of CI-like complex assembly.

### Single CI-like Structures Show Quantized NEMO Recruitment Regardless of Affinity

Our results show that CI-like complexes can form robustly in response to a range of IL-1β variants that have distinct binding characteristics. From the basis of receptor-ligand affinity and contact duration being essential contributors to signal transmission within the cell, our intuition is that IL-1β variants with lower aggregate affinity will produce lower signaling flux into the cell. One way to invoke aspects of classic receptor signaling to the IL-1β–IL-1R1, is to consider the contact duration relative to the aggregate affinity of the cytokine for the mature receptor complex. Here, the ligand-bound form of the complex will contribute positively to aspects of ubiquitin chain growth and stability through E3 ligases, leading to NEMO recruitment, whereas the unbound form does not (Figure 4A). Both bound and unbound forms are subject to degradation of the complex through the action of deubiquitinating enzymes (DUBs) as described previously^40,42,50^.

**Fig. 4:**
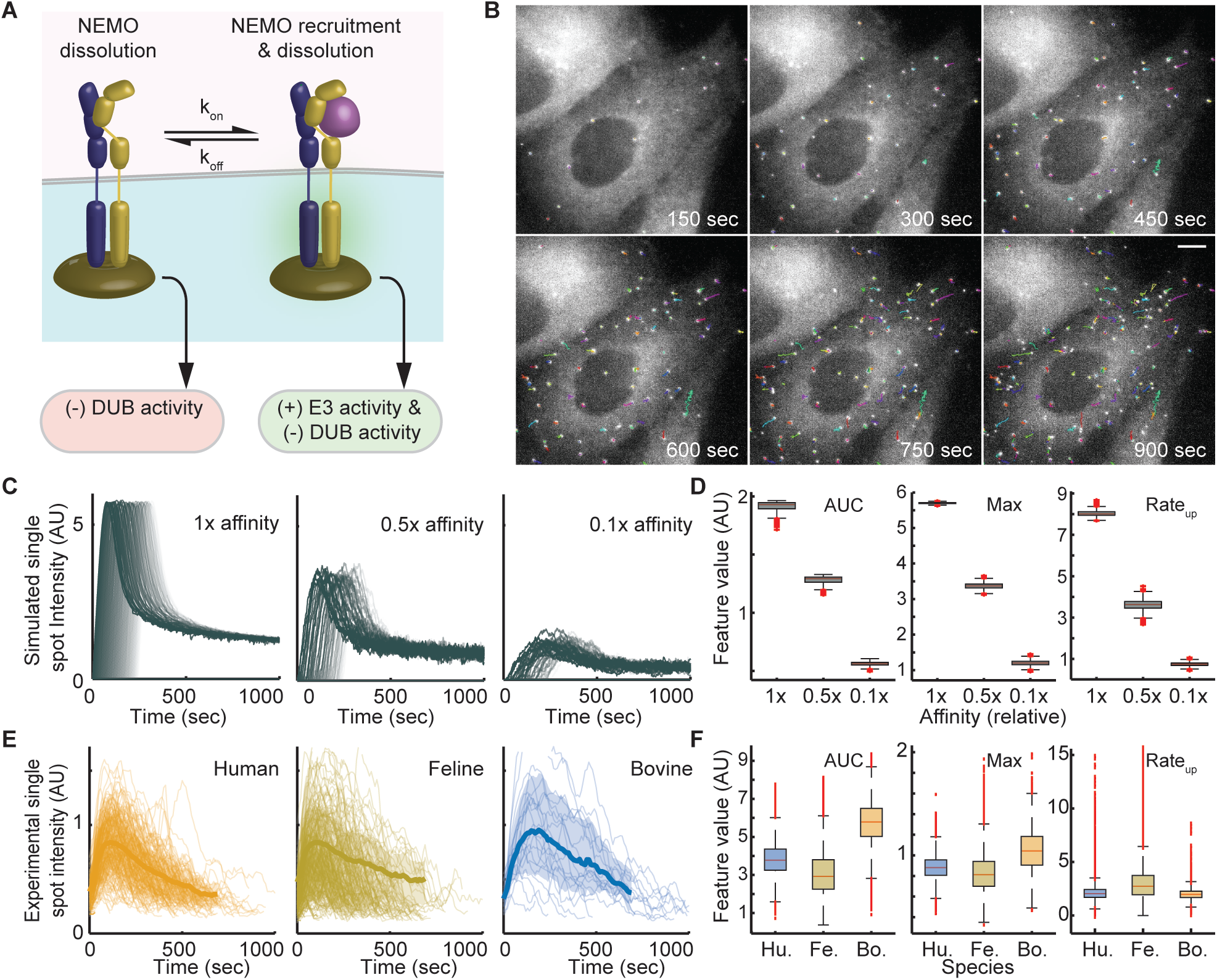
GFP-NEMO assemblies show quantized dynamics that are independent of IL-1β affinity. **A.** Schematic of classical receptor signaling demonstrating how contact duration influences behavior of single receptor complexes. In the cytokine-bound state, E3 activity promotes ubiquitin chain growth and recruitment of NEMO while DUBs contribute to ubiquitin chain disassembly. In the cytokine unbound state, NEMO complexes are degraded by basal DUB activity. **B.** Representative rapid time-lapse imaging results showing single particle tracks of EGFP-NEMO complexes in a single-cell. Scale bar, 20 µm. See also Movie S2. **C.** Stochastic simulations of spot-intensity time courses for single CI-like complexes, where affinity influences contact duration, and consequent NEMO recruitment. See Table S4 for parameter values. See also Figure S4 for alternative affinity-dependent mechanisms. **D.** Box plots of various descriptors from simulated trajectories illustrating the effects of a one-order-of-magnitude reduction in affinity on single-complex dynamics. **E.** Fast imaging and SPT of EGFP-NEMO complexes in live cells exposed to IL-1β for the indicated species. Each single trajectory represents a single assembly, with the average trajectory in bold and the standard deviation shaded. Time-courses are aggregated from multiple single cells. **F.** Box plots analogous to (B) from experimental time courses of single complexes. Despite an expectation of approximately three-orders-of-magnitude reduction in affinity for Bovine relative to Human IL-1β, single spot trends do not show significant reductions (left-tail t-test, *p* > 0.05 for all).

Through the combination of models and rapid imaging experiments coupled to SPT (Figure 4B) we tested the impact of IL-1β aggregate affinity on the dynamics of NEMO recruitment to CI-like receptor assemblies. We previously described a stochastic model simulating spot formation and dissociation events to describe the kinetics of IKK recruitment to CI assemblies^42^. While this framework does not capture the fully adaptive behavior seen experimentally, it offers a minimally sufficient model for simulating the growth and decay processes as observed for fluorescent spots. Using the model we first simulated the effects of contact duration on activating complex growth mechanisms. With the model we observed that reducing the aggregate ligand affinity: i) reduces the AUC of NEMO recruited to each simulated CI-like complex; ii) reduces the Max number of NEMO molecules per simulated complex; and, iii) reduces the rate of complex formation (Figure 4C and D). We also simulated the effect of contact duration on the rate of ubiquitin-mediated growth and NEMO recruitment, as well as the effects of contact duration on feedforward-mediated complex degradation with similar effect (Figure S4; Table S4). In all cases, simulations suggested that we should expect a significant 4- to 8-fold reduction of NEMO recruitment at CI-like structures formed in response to ligands with feline-like aggregate affinity and much greater reductions from ligands with bovine-like affinity.

Next, we used high frame-rate imaging to measure and track the intensity of EGFP-NEMO at single assemblies following exposure to human, feline, or bovine IL-1β (Figure 4B; Movie S2). To ensure that single particle tracks were directly comparable and minimally impacted by photobleaching, we only considered new spots that formed within the first 4 minutes of each imaging experiment and pooled spots from multiple cells. Remarkably, when tracked in live cells, descriptors for single complex trajectories did not show significant reductions in quantitative features (left-tail t-test, *p* > 0.05). Instead, bovine IL-1β-induced spots showed slight increases against the model-predicted trend, although these differences are likely due to the low number of single trajectories that could be tracked as described above and used for bootstrapping. Overall, trajectories of CI-like assemblies are strikingly similar, both when comparing between different cells and the response of these cells to IL-1β variants with highly dissimilar aggregate affinity. Taken together, our results show that single receptor complexes show a quantized behavior – once the complex becomes mature there is a predictable pattern of downstream signaling flux within the cell that is independent of the aggregate affinity of IL-1β for the receptor and the mature receptor complex.

## Discussion

Many rapid biological processes are mediated by affinity-dependent signaling, and a key question is how affinity impacts cytokine receptor signaling systems that can operate on larger molecular scales and longer temporal scales. Our experiments and computational analysis reveal that sequence and structure variants of IL-1β from different species are capable of binding to human IL-1R1 with a broad range of affinities, enabling us to evaluate the impact of affinity on cytokine signaling. By combining structure-based modeling with live-cell imaging experiments, we observed that species-specific IL-1β variants elicit signaling-competent CI-like assemblies observed by the formation of EGFP-NEMO puncta. When compared with human, IL-1β from other species all showed reduced sensitivity and required order(s) of magnitude higher concentrations to elicit comparable responses. The response sensitivity between non-human species stratified roughly in proportion with their predicted affinity. By contrast, observations from our models and SPT experiments further converged to demonstrate that CI-like assemblies recruit NEMO in a consistent and reproducible pattern, regardless of species and affinity.

Previously, when comparing either CI or CI-like assemblies in cells exposed to human TNF and IL-1, we observed that fluorescence properties of EGFP-NEMO spots from the same cytokine were highly similar across a broad range of concentrations^42^. Furthermore, we observed that typical TNF-induced CIs recruit approximately a third the total amount of EGFP-NEMO at each receptor complex with single-spot trajectory dynamics that were distinct when compared with IL-1β responses. CI and CI-like assemblies therefore have predictably dissimilar EGFP-NEMO recruitment dynamics.

Together with our observations here that explicitly measure the impact of cytokine affinity on CI-like spots, it suggests that cytokine receptors have evolved distinct mechanisms from the classical model. Based on these observations, we propose that IKK regulation at CI and CI-like assemblies is quantized, where the amount of signaling flux from each receptor complex is predictable and specific to the type of activated receptor complex. Through quantization, we surmise that the identity of the extracellular cytokine can be encoded in signaling space: for example, cytokine specific differences in abundance and timing of NEMO recruitment at CI and CI-like structures encodes a distinctly quantized ‘packet’ of signaling flux that is interpretated by the cell. In turn, quantized encoding can be beneficial to inflammatory responses by enabling the cell to regulate shared signaling pathways in distinct ways that encode information about different classes of extracellular messages and threats^51^.

Quantization can also have important consequences for the interpretation of inflammation and related therapeutic strategies. For example, mutations or engineered IL-1β variants that impact the cytokine-receptor affinity are likely to alter the range of cytokine concentration sensitivity by fine-tuning the activation threshold. However, through quantization, such perturbations on their own will not alter the amount and dynamics of IKK activated at each CI-like complex. It follows that quantized signaling is robust to perturbations that alter affinity because sensitivity can be corrected by upregulating or downregulating receptor expression, and each receptor complex is still producing the correct quantized response. In contrast, perturbations that alter the composition and activity of cytoplasmic constituents of CI-like assemblies, such as E3s or DUBs, may result in a confused inflammatory response that fails to encode the correct quantized signal for a particular cytokine receptor. Therefore, inflammatory diseases and related therapeutics may benefit from being evaluated under the lens of quantized signaling.

Our comparative modeling of IL-1β–IL-1R1 complexes across species reveals nontrivial variation in predicted binding affinity, reflecting a likely divergence in cross-species cytokine–receptor interactions. While our simulations focus on the primary ligand–receptor interface, it is important to emphasize that this is only one step in the assembly of the mature CI-like signaling complex. Crucially, the recruitment of IL-1R3 is dependent not only on IL-1β’s affinity for IL-1R1 but also on ligand-induced conformation changes in the receptor, as well as the orientation of transmembrane domains and cytoplasmic components. Since there is no ground truth information for many of these structures, our analysis focused on the extracellular domain interaction, which represents the most conserved and structurally tractable portion of the complex. Although useful for comparative purposes and leading to results that generally agree with experimentally defined “aggregate affinities”, a limitation of our study is that the predicted binding conformations are not guaranteed to recapitulate binding poses *in vivo*. Moreover, the relatively short simulations, while capturing trends, limit our ability to make precise quantitative comparisons, particularly in systems where long-timescale conformational sampling may influence the energy landscape. Despite these limitations, the differences in predicted binding free energy suggest the likelihood for competition between IL-1β variants across species for related receptor pools, as well as the decoy receptor IL-1R2.

While distinct from the classical model, quantized receptor signaling still relies on affinity and contact duration to establish the sensitivity threshold. In terms of quantized signaling, it remains to be established whether other non-cytokine receptors also have similar mechanisms, and how cells use quantized signaling to productively encode information. Conceivably, benefits to the cell from quantized signaling can be multi-fold. E.g., contributing to mechanistic robustness, parsimonious and predictable amplification of extracellular signals, or possibly contributing to time-domain communication strategies akin to Morse code, where messages are conveyed through timing and duration rather than amplitude alone. As future studies further characterize the mechanisms of dynamics and quantization of CI-like assemblies, we will gain more understanding of underlying biological design principles and identify new control points that shape cytokine responses.

## Author contributions

R.E.C.L., A.H.K., and B.K. conceived of the study. A.H.K., B.K., and J.Y. performed the analysis of live-cell images. A.H.K., B.K., and R.E.C.L. developed and interpreted the dynamic systems models. B.K. and D.R.K. developed and interpreted the structural models. R.E.C.L., A.H.K., and B.K. wrote the manuscript.

## Methods

### Cell Culture

U2OS cells endogenously expressing EGFP-NEMO and mCherry-RelA¹ were cultured in McCoy’s 5A medium (Thermo Fisher) supplemented with 10% fetal bovine serum (FBS; Corning), 100 U/mL penicillin, 100 µg/mL streptomycin, and 0.2 mM L-glutamine (Invitrogen). Cells were maintained at 37°C in a humidified incubator with 5% CO_2_ and routinely tested for mycoplasma contamination.

### Recombinant Cytokines

Recombinant IL-1β proteins were obtained from multiple commercial sources for cross-species stimulation assays. Human, mouse, and rat IL-1β recombinant proteins were purchased from ThermoFisher (PeproTech brand; catalog numbers: 200-01B-2UG, 211-11B-2UG, and 400-01B-2UG, respectively). Bovine IL-1β was sourced from ThermoFisher (Invitrogen; RBOIL1BI). Feline and rabbit IL-1β/IL-1F2 proteins were acquired from R&D Systems (catalog numbers: 1796-FL-010 and 7464-RB-010, respectively). All cytokines were reconstituted and aliquoted according to the manufacturers’ instructions and stored at –80°C until use.

### Live-Cell Imaging

Live-cell imaging was performed in an environmentally controlled chamber (37°C, 5% CO₂) using a DeltaVision Elite microscope (GE Healthcare) equipped with a pco.edge sCMOS camera and an Insight solid-state illumination module. U2OS cells expressing fluorescent protein–tagged RelA were seeded at a density of 8,000 cells per well in no. 1.5 glass-bottom 96-well imaging plates (Matriplate) 24 hours prior to imaging. One hour before acquisition, culture medium was replaced with phenol red–free FluoBrite DMEM (Gibco, A18967-01) supplemented with 10% fetal bovine serum (FBS; Corning), 100 U/mL penicillin, 100 µg/mL streptomycin, and 0.2 mM L-glutamine (Invitrogen).

EGFP-NEMO images were acquired every 3 minutes for 6 hours using FITC filter sets and a 60× LUCPLFLN oil immersion objective (Movie S1). Z-stacks were collected with eight planes at 0.5 µm intervals using a 0.04-second exposure time and 32% transmission. To image single-complex dynamics, EGFP-NEMO images were acquired every 10 seconds for 1 hour using the same settings as above (Movie S2). For NEMO spot tracking experiments, cells were stimulated with the indicated concentrations of IL-1 immediately before imaging. For imaging single-complex dynamics, a concentration of 10 ng/mL was used for IL-1β from human and feline, while a concentration of 500 ng/mL was used for bovine.

### Detection and Quantification of EGFP-NEMO spots

EGFP-NEMO puncta were detected and quantified using dNEMO, a custom computational tool optimized for analyzing fluorescent puncta in fixed and live-cell time-lapse images^42,52^. A user-defined detection threshold ranging from 1.5 to 2.8 was applied uniformly across all datasets. Puncta were considered valid only if they appeared in at least two contiguous slices within the 3D image stack (eight slices total). Pixel intensities for each punctum were background-corrected by averaging values from an annular ring (1-pixel width and offset) surrounding the spot. Single cells were manually segmented using dNEMO’s keyframing function. Spot features were extracted for each newly formed punctum following stimulation, yielding time-resolved, single-cell measurements.

### Single Particle Tracking of EGFP-NEMO spots

EGFP-NEMO spot location and intensity data obtained from dNEMO were used for single-spot tracking with the *uTrack* package in MATLAB^53,54^. Tracking parameters were optimized to improve performance for IL-1–induced puncta. Specifically, the maximum allowed gap between detections was reduced from five to three frames, and the gap-closing penalty was lowered from 1.5 to 1. The minimum track segment length used for gap closing was increased from one to three frames. Spot properties, including intensity and size, were associated with their corresponding trajectories to compute integrated intensity over time. To minimize the effects of photobleaching, only spots that formed within the first four minutes of imaging were included in the analysis.

### Bootstrapping of Single-Complex Spot Trajectories

To account for the low number of EGFP-NEMO spots detected in U2OS cells stimulated with 500 ng/mL Bovine IL-1β, bootstrapping was performed to enable statistical comparisons across conditions. For each IL-1β species condition (Human, Feline, and Bovine), we resampled the set of tracked EGFP-NEMO trajectories 100,000 times using a sample size of 5 per iteration. Resampling was implemented using a custom MATLAB script.

### Extracting Features from Spot Trajectories

Quantitative descriptors of EGFP-NEMO trajectories were extracted using custom MATLAB and Python scripts from either single-cell or single-complex data. For long time-course experiments, trajectories were generated based on the number of NEMO puncta per cell over time. For short-term tracking experiments, trajectories reflected the integrated intensity of all NEMO spots within a cell at each time point. The following metrics were computed for each trajectory:

1. Area under the curve (AUC): Total integrated value of the trajectory, representing either the cumulative number of NEMO spots or the total spot intensity per cell.
2. Maximum value (Max): Peak number of NEMO spots or peak integrated intensity.
3. Rate of rise (*Rate_up_*): Maximum slope (absolute value) calculated from a linear fit to three consecutive time points between time zero and the time of maximum signal.
4. Rate of decay (*Rate_down_*): Maximum slope (absolute value) from a linear fit to three time points between the time of maximum signal and the end of the trajectory.
5. Time of maximum (*t_max_*): Time point at which the trajectory reaches its maximum value.
6. Time to half-maximum rise (*t_50,up_*): Time at which the trajectory first reaches 50% of its maximum value.
7. Time to half-maximum decay (*t_50,down_*): Time at which the trajectory falls back to 50% of its maximum value.
8. Full width at half maximum (FWHM): Duration between *t_50,up_* and *t_50,down_*, indicating the width of the response at half-maximum intensity.
9. Adaptation time: Time at which the trajectory first reaches its lower plateau value.

### Sigmoidal Fitting of AUC and Max Features

Sigmoidal curve fitting of extracted AUC and Max features from NEMO spot number trajectories was performed using a custom Python script. For each species and cytokine concentration, the mean feature value was calculated and used as the input for curve fitting. A standard sigmoid equation was applied:

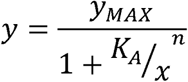

Fits were computed using the curve_fit function from the *SciPy* package. Estimated parameter values and corresponding R² scores are summarized in Table S1.

### Quantifying Noise in Feature Distributions Using the Fano Factor

To evaluate cell-to-cell variability in extracted features from NEMO spot number trajectories, we computed the Fano factor, a common measure of noise defined as the variance-to-mean ratio. For each species and cytokine concentration, we extracted single-cell values for five dynamic descriptors: AUC, Max, T_MAX_, FWHM, and Adaptation Time.

Using custom Python scripts, we first grouped single cell feature values by species and stimulation dose. For each group, we calculated the mean and variance, then computed the Fano factor as:

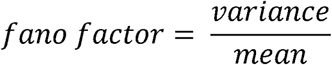

Values were excluded if the group mean was zero or undefined. To visualize how noise varied across species and concentrations, we generated two complementary plots per feature: (1) a line plot of the Fano factor across cytokine concentrations for each species, and (2) a species-wise box plot showing the distribution of Fano values across all doses.

This analysis allowed us to quantify the extent of single-cell heterogeneity for each extracted feature, across both species and stimulation conditions.

### Stochastic Simulation of CI-Like Complex Formation Dependent on Binding Affinity

We modelled the bulk assembly of CI-like complexes using the Gillespie algorithm. This model was designed to determine the range in relative IL-1β–IL-1R1 affinity necessary to recapitulate experimental measurements of EGFP-NEMO puncta numbers and timing. The key variable simulated was the number *N* of mature CI-like complexes resulting from the assembly cascade previously described. The relative binding affinity *Aff* altered only the on-rate *K_on_* and off-rate *K_off_* of IL-1β–IL-1R1 binding, where *K_on_* = *K_on_*_,0_ * *Aff* and *K_off_* = *K_off_*_,0_ / *Aff*. All other parameters were held constant. The values for each parameter are described in Table S2.

### Sigmoidal Fitting of Simulated AUC and Max Features

Sigmoidal curve fitting of extracted AUC and Max features from stochastically simulated NEMO spot number trajectories was performed using a custom Python script. For each relative affinity and cytokine concentration, the mean feature value was calculated and used as the input for curve fitting. A standard sigmoid equation was applied:

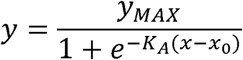

Fits were computed using the curve_fit function from the *SciPy* package. Estimated parameter values and corresponding R² scores are summarized in Table S3.

### Computational Model (HyDeS) to Simulate Single-Complex Dynamics Dependent Upon Contact Duration

We modelled the intensity dynamics of individual EGFP-NEMO puncta using a hybrid deterministic–stochastic (HyDeS) framework. This model was designed to test how ligand–receptor contact duration influences the assembly of signalosome-like structures. Two key variables were simulated: the intensity *I* of each NEMO spot, representing a continuous approximation of ubiquitin chain size and NEMO activity at that site; and a feedback variable *x_basal_*, which serves as a proxy for the influence of basal deubiquitinating enzyme (DUB) activity present in resting cells. Basal feedback was modeled to increase proportionally with spot intensity, representing DUB recruitment to active complexes.

The model comprises *2N* coupled ordinary differential equations (ODEs) for *N* NEMO spots: one equation for the intensity *I_i_* of each spot and one for the corresponding feedback variable 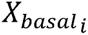. Simulations were performed with 75 individual spots forming at regular intervals, approximating the response of a single cell following cytokine stimulation.

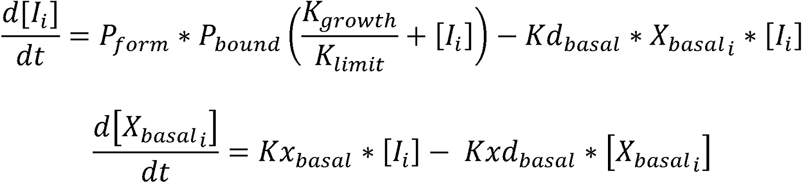

The variables and parameter values used in the model are as described in tables S2. We examined the effects of contact duration on the single complex dynamics in three different ways: (1) toggling complex formation on and off based on binding state, corresponding to lower *p_bound_* for lower affinities; (2) modulating the rate of complex formation upon binding, corresponding to lower *K_growth_* for lower affinities; and (3) suppressing DUB activity at higher affinities. For the latter, *Kd_basal_* is divided by the affinity, which reflects DUB inhibition due to higher affinities, in the following way:

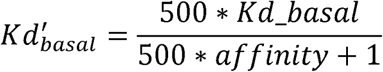

An arbitrary multiplier was used to scale up the effects of affinity on *Kd_basal_*. The descriptions and default values for all parameters are listed in Table S4.

### Structurally Informed Distance Tree Construction

Multiple sequence alignment via MAFFT was performed on recombinant IL-1β sequences from each species^55^. Human IL-1β binding regions were annotated using a distance threshold of 4.5Å between residues of IL-1β to IL-1R1 or IL-1R3 in the complex’s crystal structure (PDB: 4DEP). Unweighted Pair-Group Method using Arithmetic averages trees were constructed with BLOSUM62 similarity for both the full sequence and annotated binding region.

### Chimeric Structure Prediction

Predicted structures for each IL-1β–IL-1R1 complex were generated using the Alphafold3 server, which was provided the recombinant species-specific IL-1β sequence and the human IL-1R1 sequence. All five highest-pLDDT predictions were considered as optimal binding candidates.

### Molecular Dynamics System Preparation

The crystal structure for the human IL-1β–IL-1R1 complex (PDB: 1ITB) was downloaded, prepared for use in the AMBER suite with PDBfixer v. 1.11, parameterized with the AMBER ff15IPQ force field^56^, and solvated with a 12Å TIP3P water box with 12Å padding.

For each chimeric system, Open Babel v. 3.1.0^57^ was used to convert AF3 structures to a complexed IL-1β–IL-1R1, isolated IL-1β, and isolated IL-1R1 PDB file for each of the 5 predicted structures. Each was then prepared for use in the AMBER suite via pdb4amber v. 1.6.dev.^58^ and parameterized with the AMBER ff15IPQ force field. An additional neutralized and 12Å TIP3P solvated structure file was produced for each complex.

### Molecular Dynamics Simulation

Classical MD simulations for each solvated complex were performed with the Amber 24 software package^58^ with a 2-fs time step. Water and ions were first minimized for 50ps with a protein restraint weight of 2.0 kCal/mol Å^2^. Under the same protein restraint, each system was heated with constant volume from 100K to 298K over 50 ps, followed by 50 ps of constant pressure relaxation. Each system was then equilibrated under no restraint for 1 ns. Finally, five successive 5ns production simulations for each system with no restraint were performed. Velocities were reset between each simulation.

### Free energy Estimation with MM/GBSA and MM/PBSA

100 frames from each of the five 5ns simulations were used in MM/GBSA and MM/PBSA^46^ to estimate the total binding free energy and per-residue decomposition of binding free energy respectively. The lowest MM/GBSA binding free energy simulation was chosen as the ideal conformation for MM/PBSA and structural comparison.

## Acknowledgements

We thank other members of the Lee lab and colleagues in the Department of Computational and Systems Biology for many helpful discussions. This work was supported by generous funding to R.E.C.L. from NIH grant R35GM119462, and NIH grant R35GM140753 to D.R.K. J.Y. received support from the ARCS Foundation Pittsburgh Chapter.

## Supplementary figure legends

**Supplementary Figure 1: Comparative Structural and Energetic Analysis of IL-1**β**– IL-1R1 Interactions Across Species**

**A.** Multiple sequence alignment of IL-1β sequences from a panel of 12 species. Predicted residues involved in IL-1R1/IL-1R3 binding denoted by red rectangles. **B.** Differences between the crystal structure and predicted structure for the IL-1β–IL-1R1 complex in terms of predicted binding conformations and contributions to changes in free energy resulting from binding. **C.** Backbone RMSD throughout simulation for the 6 predicted IL-1β–IL-1R1 complexes. **D.** Estimated changes in binding free energy of generated IL-1β/I-L1R1 complexes relative to that of the human crystal structure.

**Supplementary Figure 2: Cross-Species Comparison of EGFP-NEMO Complex Formation and Intensity Profiles in Response to IL-1β Stimulation**

**A.** Time-lapse images of EGFP-NEMO in live U2OS cells stimulated with 250 ng/mL IL-1β from the indicated species. Line scans were performed at 24 minutes post-stimulation (blue lines). Scale bar, 20 µm.

**B.** Line intensity profiles of two representative EGFP-NEMO spots corresponding to line scans shown in (A), from cells stimulated with 250 ng/mL IL-1β from the indicated species.

**Supplementary Figure 3: Experimental and Simulated Dose–Response Analysis of EGFP-NEMO Complex Formation Using the MAX Descriptor.**

**A.** Sigmoid curves fitted to the mean maximum spot number (MAX) from experimental single-cell time courses across IL-1β concentrations and species, revealing species-specific dose–response relationships. The AC10, defined as the concentration at which 10% of the maximal response is reached, is indicated. See Table S1 for fit parameters.

**B.** AC10 values for each species derived from the MAX descriptor fits shown in (A), indicating the concentration at which 10% of the maximal response is reached.

**C.** Stochastic simulations of spot number time courses across species and concentrations, using affinity parameters optimized to match experimental data. Fitted affinity values are shown above each plot, representing relative signaling competency of each species. See also Table S2. **D**. Simulated dose–response curves for the MAX descriptor, recapitulating relative IL-1β–IL-1R1 affinity-dependent complex formation dynamics. See also Table S3. **E**. Simulated AC10 values derived from the MAX descriptor for a range of IL-1β–IL-1R1 affinities, reflecting predicted sensitivity to IL-1β across species.

**Supplementary Figure 4: Simulated Effects of Contact Duration on EGFP-NEMO Single-Complex Dynamics.**

**A.** Conceptual schematic extending Figure 4A, illustrating three hypothetical mechanisms by which ligand–receptor contact duration may influence EGFP-NEMO complex dynamics: (1) toggling complex formation on and off based on binding state; (2) modulating the rate of complex formation upon binding; and (3) suppressing DUB activity at higher affinities. **B.** Stochastic simulations of single-complex spot intensity trajectories under each of the three mechanisms described in (A). See Table S2 for parameter values. Box plots summarize key trajectory descriptors, demonstrating how a one-order-of-magnitude reduction in receptor–ligand affinity alters single-complex dynamics.

