## Supplementary figures and images for "NEMO recruitment at single cytokine-receptor complexes shows quantized dynamics independent of ligand affinity"

### Supplemental Figure 1

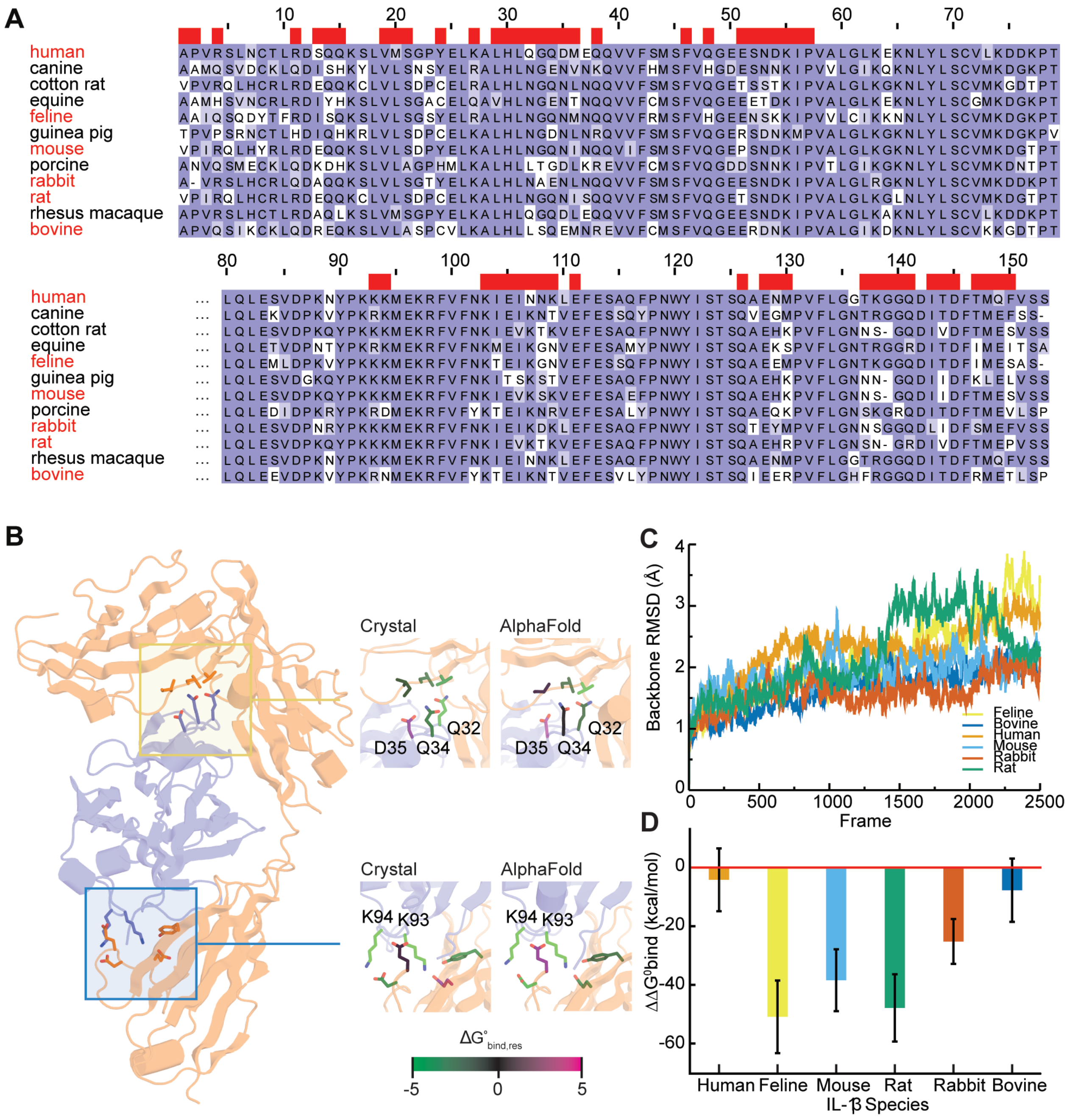

### Supplemental Figure 2

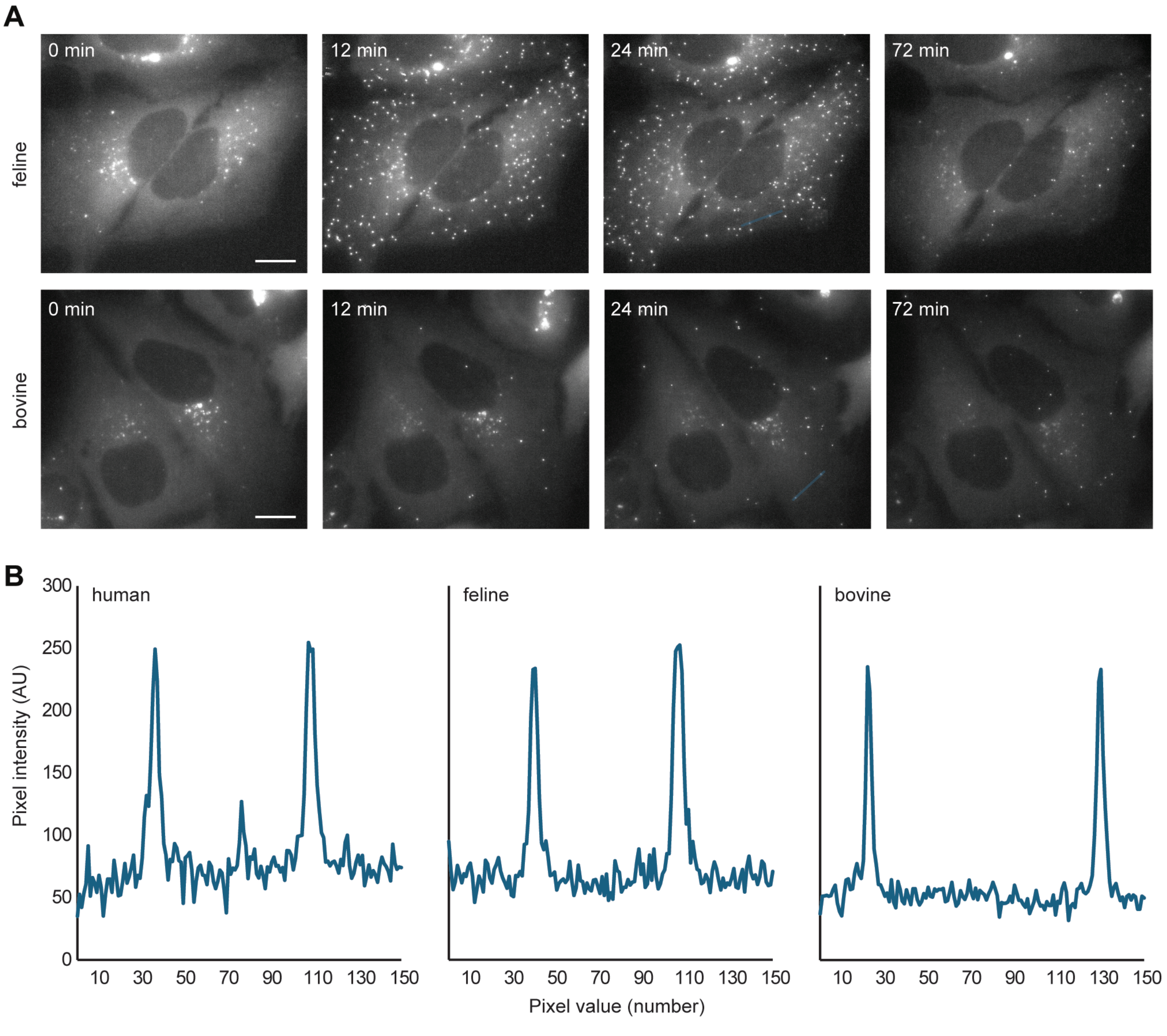

### Supplemental Figure 3

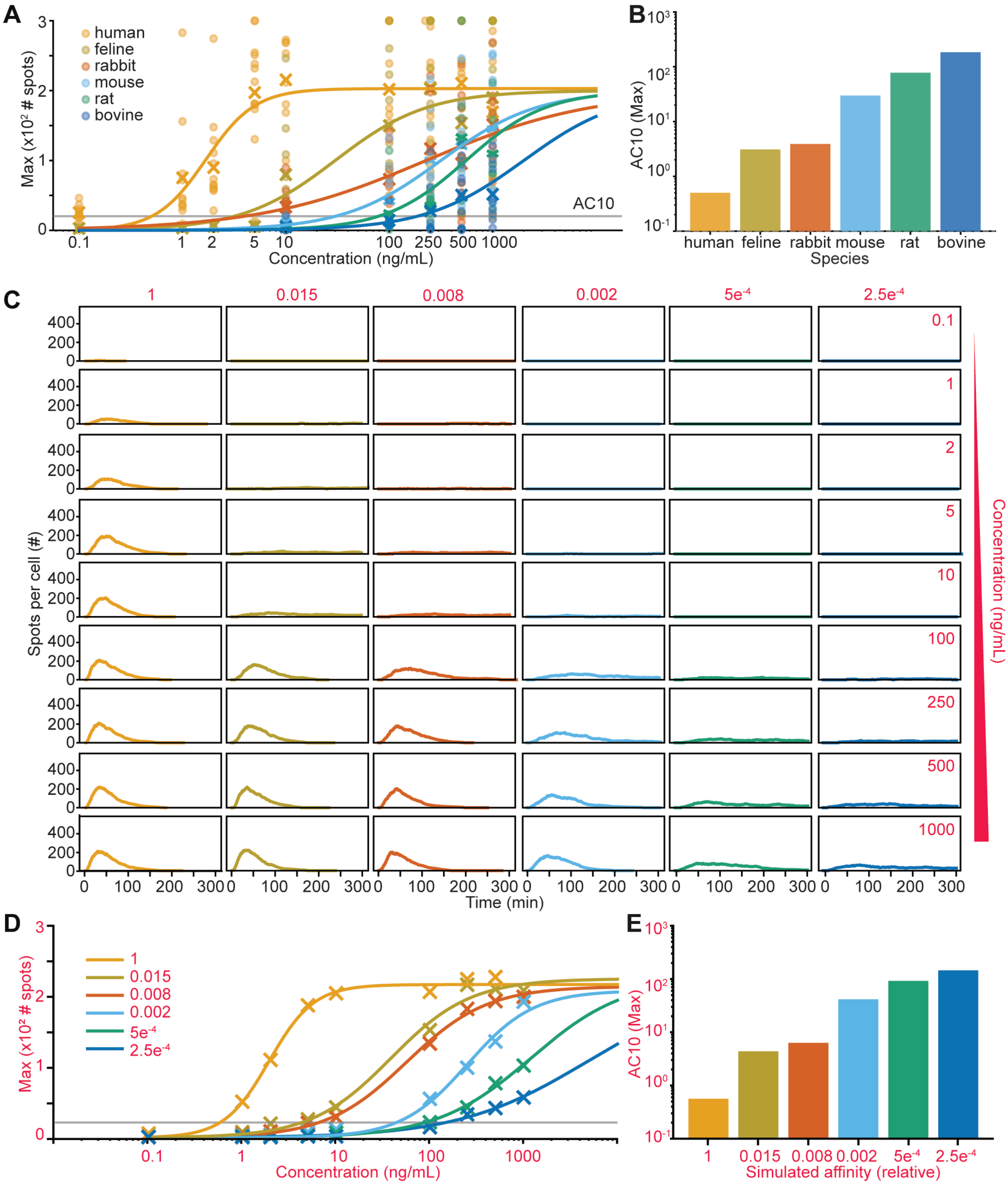

### Supplemental Figure 4

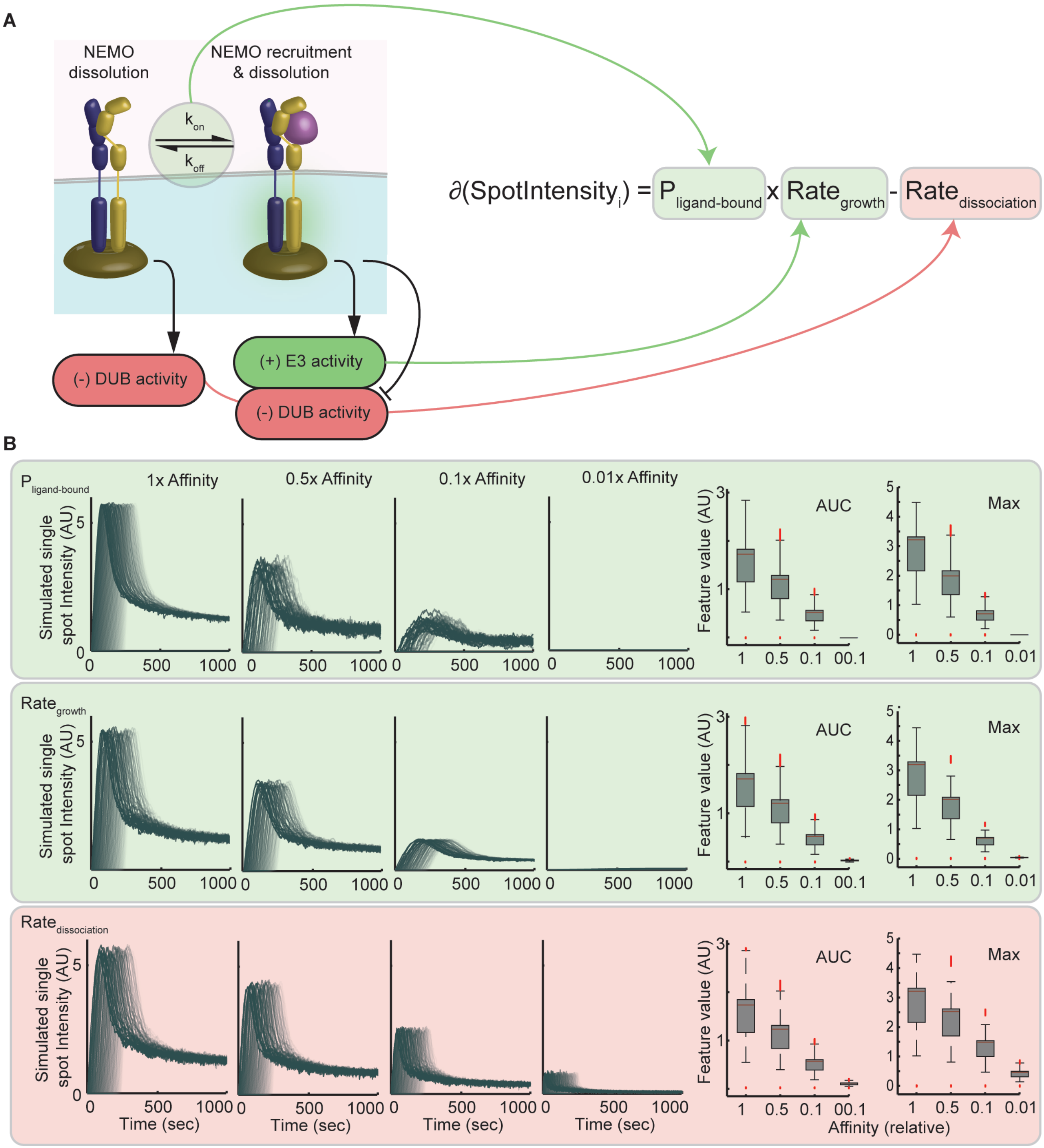
