## Supplemental Table 1 for "NEMO recruitment at single cytokine-receptor complexes shows quantized dynamics independent of ligand affinity"

**Table S1.** Parameters from sigmoidal fits to AUC and MAX descriptors of EGFP-NEMO spot number timecourses taken from live cell imaging data.

|  | AUC | | | | | MAX | | | | |
| --- | --- | --- | --- | --- | --- | --- | --- | --- | --- | --- |
| Species | y_MAX_ | K_A_ | n | AC_10_ | R^2^ | y_MAX_ | K_A_ | n | AC_10_ | R^2^ |
| Human | 2900 | 1.8 | 1.4537 | 0.4 | 0.831 | 203 | 1.7 | 1.8217 | 0.5 | 0.911 |
| Feline | 3000 | 39.6 | 0.9012 | 3.5 | 0.959 | 200 | 30.0 | 0.9720 | 3.1 | 0.945 |
| Rabbit | 3000 | 136.5 | 0.7222 | 6.5 | 0.605 | 200 | 227.5 | 0.5395 | 3.9 | 0.867 |
| Mouse | 2900 | 206.6 | 0.9839 | 22.1 | 0.917 | 200 | 315.6 | 0.9315 | 29.8 | 0.981 |
| Rat | 3000 | 375.0 | 1.9000 | 118.0 | 0.794 | 200 | 553.7 | 1.1199 | 77.8 | 0.806 |
| Bovine | 3000 | 1200.0 | 1.1386 | 174.2 | 0.882 | 200 | 2000.0 | 0.9217 | 184.4 | 0.787 |
