## Supplemental Table 2 for "NEMO recruitment at single cytokine-receptor complexes shows quantized dynamics independent of ligand affinity"

**Table S2.** Parameter descriptions and default values used for stochastic simulations run for EGFP-NEMO spot number timecourses.

| **Parameter** | **Description** | **Default Value** |
| --- | --- | --- |
| K_1,on_ | Rate of IL-1B binding to IL1R1, linearly dependent on their relative binding affinity. | 1.8*10^-4^ |
| K_1,off_ | Rate of IL-1B unbinding to IL1R1, inversely linearly dependent on their relative binding affinity. | 3*10^-6^ |
| K_2,on_ | Rate of IL1R3 binding to formed IL-1B/IL1R1 complex | 9*10^-5^ |
| K_2,on_ | Rate of IL1R3 unbinding to formed IL-1B/IL1R1 complex | 3*10^-6^ |
| K_3,on_ | Rate of Myd88 binding to the cytoplasmic domains of the assembled receptor complex to initiate NEMO recruitment. | 6*10^-8^ |
| K_3,off_ | Rate of Myd88 unbinding to the cytoplasmic domains of the assembled receptor complex. | 3*10^-9^ |
| K_i_ | Rate of internalization or degradation of the complex. | 0.036 |
