## Supplemental Table 3 for "NEMO recruitment at single cytokine-receptor complexes shows quantized dynamics independent of ligand affinity"

**Table S3.** Parameters from sigmoidal fits to AUC and MAX descriptors of EGFP-NEMO spot number timecourses taken from stochastic simulations.

|  | AUC | | | | | MAX | | | | |
| --- | --- | --- | --- | --- | --- | --- | --- | --- | --- | --- |
| Affinity | y_MAX_ | K_A_ | X_0_ | AC_10_ | R^2^ | y_MAX_ | K_A_ | X_0_ | AC_10_ | R^2^ |
| 1 | 3000 | 0.2 | 4.9691 | 0.6 | 0.988 | 216 | 0.3 | 4.1989 | 0.6 | 0.988 |
| 0.015 | 2950 | 1.0 | 3.0028 | 1.7 | 0.992 | 225 | 1.6 | 2.3544 | 4.5 | 0.992 |
| 0.008 | 3040 | 1.1 | 4.1901 | 3.8 | 0.987 | 214 | 1.8 | 2.3370 | 6.5 | 0.999 |
| 0.002 | 3041 | 1.8 | 3.1574 | 11.7 | 0.999 | 208 | 2.4 | 2.8445 | 42.5 | 0.991 |
| 0.0005 | 2958 | 2.3 | 3.3552 | 40.8 | 0.999 | 225 | 3.1 | 1.9889 | 93.9 | 0.999 |
| 0.00025 | 3047 | 2.6 | 2.8722 | 69.2 | 0.999 | 217 | 3.7 | 1.4503 | 148.2 | 0.999 |
