## Supplemental Table 4 for "NEMO recruitment at single cytokine-receptor complexes shows quantized dynamics independent of ligand affinity"

**Table S4.** Parameter descriptions and default values for HyDeS model, which simulates EGFP-NEMO single complex spot intensity timecourses.

| **Parameter** | **Description** | **Default Value** |
| --- | --- | --- |
| P_form_ | Probability of spot formation dependent on time. A formed spot is assumed to stay active. | Uniform distribution of spots forming at regular intervals across given time scale. |
| P_bound_ | Probability of ligand remaining bound to receptor, driving recruitment of NEMO. | 0.9 |
| K_growth_ | Intrinsic Michaelis-Menten like rate of ubiquitin polymerization. | 1.12*10^-7^ |
| K_limit_ | Michaelis-Menten like kinetic constant limiting intrinsic polymerization as spot intensity increases. | 0.1 |
| Kd_basal_ | Rate of feedback activity on individual spots from basal DUB levels. | 0.032 |
| Kx_basal_ | Rate of activation of feedback due to ubiquitin polymerization and basal DUB recruitment for each spot. | 23.3 |
| Kxd_basal_ | Decay rate of basal feedback. | 8.7*10^-4^ |
